# A general kernel machine comparative analysis framework for randomized block designs

**DOI:** 10.1101/2025.02.06.636977

**Authors:** Hyunwook Koh

## Abstract

**Motivation:** There are numerous potential confounders, including genetic, environmental, technical, and demographic factors. These factors may be known or unknown, measured or unmeasured; hence, it is extremely challenging to capture them in downstream data analysis. However, randomized block design is an efficient design technique to control confounding factors and to reduce variability within subjects. This helps prevent spurious discoveries and boost test power. I also note that kernel machine comparative analysis is widely employed in high-dimensional omics studies to boost test power by combining possibly weak effects from multiple underlying variants, and also to explore various linear or nonlinear patterns of disparity.

**Results:** In this paper, I introduce a general kernel machine comparative analysis framework for randomized block designs, named as KernRBD, to investigate the effects of treatments (e.g., medical treatment, environmental exposure) on the underlying variants. KernRBD is unique in its range of functionalities, including the computation of *P*-value for global testing and adjusted *P*-values for pairwise comparisons, as well as visual representation through ordination plotting. KernRBD is practical, requiring only a kernel as input, and also robustly valid based on a resampling scheme not requiring the assumption of normality to be satisfied. I also introduce its omnibus test for a unified and powerful significance testing across multiple input kernels. While its applications should be much broader, I illustrate its use through human microbiome *β*-diversity analysis *in praxis*, and its outperformance in significance testing through simulation experiments *in silico*.

**Availability and Implementation:** KernRBD is available at https://github.com/hk1785/kernrbd.

## 1 Introduction

Recent advances in high-throughput sequencing technologies have enabled faster, more affordable, and more precise quantification of various omics profiles. Consequently, researchers have been able to identify numerous biomarkers associated with the effects of treatments, demographics, environmental factors, technical factors, behavioral factors, and so on. However, this can also underscore a critical challenge for new research: the presence of potential confounding factors that can lead to spurious findings.

It is extremely challenging to control various known or unknown, measured or unmeasured confounding factors in downstream data analysis. An efficient and practical solution is to use randomized blocks from the experimental design phase, where subjects are typically the blocks of multiple observations for randomly assigned treatments [1]. Here, the use of blocks helps maintain the levels of all other unwanted factors, except for the treatment(s) of interest, consistent within each subject, thereby reducing within-subject variability [1]. The use of randomized treatment assignments can also eliminate the effect of treatment ordering, which is alternatively addressed in practice through a washout period between treatments or a pre- and post-treatment design [1]. Though researchers can also be interested in the cumulative effects of repeated treatments themselves, without the need for randomized treatment assignments. In this context, the randomized block design helps prevent spurious findings and enhance test power.

Notably, kernel machine comparative analysis is also widely employed in high-dimensional omics studies to enhance test power by combining potentially weak effects from multiple underlying genetic or microbial features [2, 3, 4]. Here, a kernel is defined as a positive semi-definite matrix that represents the similarity or relatedness for each pair of observations, which compresses high-dimensional hundreds or thousands of features. For example, the simplest linear kernel is the inner product matrix of the original feature matrix, and it is especially useful to survey the linear patterns of disparity between treatments. On the contrary, polynomial kernels are useful to survey the nonlinear patterns of disparity between treatments. More generally, a kernel is an inner product matrix of any linearly or nonlinearly transformed feature matrix. There are also various measures of similarity or relatedness, leading to various kernels to exist in practice. This enables flexible exploration of various linear or nonlinear patterns of disparity using various kernels.

In this paper, I introduce a general kernel machine comparative analysis framework for randomized block designs, named as KernRBD, to investigate the effects of treatments (e.g., medical treatment, environmental exposure) on the underlying genetic or microbial features. KernRBD is unique in its range of functionalities, including the computation of *P*-value for global testing and adjusted *P*-values for pairwise comparisons, and the visual representation using an ordination plot in a reduced coordinate space. Mercer’s theorem [5] ensures that even if the underlying function for the linear or nonlinear feature transformation is unknown, its associated reproducing kernel Hilbert space (RKHS) is well-defined [5]. This enables the practical implementation of KernRBD, requiring only a kernel as input without the need to know of its underlying real features [6]. KernRBD is also robustly valid based on a resampling scheme, not requiring the assumption of normality to be satisfied. I also note that the result can dramatically vary depending on the choice of kernel, which makes consistent interpretation challenging [2, 3, 4, 7, 6]. Therefore, I also introduce the omnibus test of KernRBD based on the minimum *P*-value test statistic [8] for a unified and powerful significance testing across multiple input kernels.

I illustrate the use of KernRBD through human microbiome *β*-diversity analysis [9, 10, 4, 11] for the following machinery and reasons. I utilize ecological kernels, including Jaccard [12], Bray-Curtis [13], unweighted UniFrac [14], weighted UniFrac [15] and generalized UniFrac [16] kernels, that represent ecological similarity for each pair of observations from different perspectives. Prior studies [4, 7, 6] have extensively examined their unique and well-distinguished performances in significance testing. All these ecological kernels are constructed via complex feature transformations, such as using presence/absence information or the full abundance spectrum, and either incorporating or ignoring phylogenetic tree information; yet, the exact forms of their underlying transformation functions are hard to retrieve [6]. Therefore, they can only be utilized within a general framework that accepts a kernel directly as input. Besides, microbiome data are highly skewed with excessive rare features and a long tail of zeros. Therefore, statistical inference based on the assumption of normality is not also valid.

A notable existing method of its kind is a distance-based analysis, known as permutational multivariate analysis of variance (PERMANOVA) [9, 10, 11], using within-block permutations. In human microbiome studies, PERMANOVA [9, 10, 11] is widely employed because of its powerful performance leveraging ecological distances that represent ecological dissimilarity for each pair of observations from different perspectives [12, 13, 14, 15, 16], as well as its nice visual representation through multidimensional scaling [17, 18, 19] for better interpretability. However, PERMANOVA [9, 10, 11] is designed only for global testing, not providing adjusted *P*-values for pairwise comparisons. Furthermore, PERMANOVA [9, 10, 11] can process ecological distances individually with no omnibus testing capabilities. Coincidentally, through simulation experiments, I also found that KernRBD is more powerful than PERMANOVA [9, 10, 11], even when using the same kernel/distance for global testing.

The remainder of the paper is structured as follows. In the *Materials and Methods* section, I detail the methodology of KernRBD. Then, in the *Results* section, I illustrate the outperformance of KernRBD through simulation experiments, as well as its application to real gut microbiome data on the effect of periodically restricted feeding (PRF) [20]. Finally, in the *Discussion* section, I provide a summary and discuss potential extensions and applications.

## 2 Materials and Methods

### 2.1 Notations and Models

Suppose that there are *n* subjects (*i* = 1, …, *n*); *m* treatments (e.g., pre-treatment vs. post-treatment(s), reference vs. comparison(s)) per subject (*j* = 1, …, *m*) with *N* = *nm* to be the total number of observations; and *p* features (e.g., genetic or microbial variants) (*k* = 1, …, *p*) possibly with *p* ≫ N. Then, let *Y*_*ij*_ denote a *p* × 1 vector of the features for the *i*-th subject and *j*-th treatment: *Y*_*ij*_ = (*Y*_*ij*1_, …, *Y*_*ijp*_)^*T*^. For simplicity, I assume one observation per treatment in each subject, but its extension to multiple observations in an unbalanced or incomplete design is easy. Now,

I formulate an effects model as in Eq. (1), or equivalently a means model as in Eq. (2).

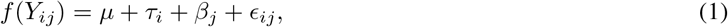

and

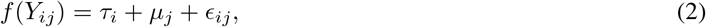

where *f* (·) is a function for a linear or nonlinear transformation; *μ* is a *p* × 1 vector of the overall means: *μ* = (*μ*_1_, …, *μ*_*p*_)^*T*^; *τ*_*i*_ is a *p* × 1 vector of the *i*-th subject’s effects: *τ*_*i*_ = (*τ*_*i*1_, …, *τ*_*ip*_)^*T*^; *β*_*j*_ is a *p* × 1 vector of the *j*-th treatment’s effects: *β*_*j*_ = (*β*_*j*1_, …, *β*_*jp*_)^*T*^; *μ*_*j*_ is a *p* × 1 vector of the *j*-th treatment’s means: *μ*_*j*_ = *μ* + *β*_*j*_ = (*μ*_*j*1_, …, *μ*_*jp*_)^*T*^; and *ϵ*_*ij*_ is a *p* × 1 vector of the independently and identically distributed errors with mean zero and variance *σ*^2^.

First, I consider global testing with the null hypothesis of no effect for any treatment as in Eq. (3), or equivalently the null hypothesis of no difference in mean for any pair of treatments as in Eq. (4).

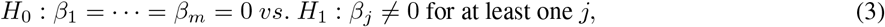

and

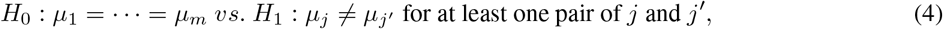

where *j* = 1, …, *m* and *j*′ = 1, …, *m* with *j* ≠ *j*′.

Second, I consider pairwise comparisons with the null hypothesis of no effect for each treatment as in Eq. (5), or equivalently the null hypothesis of no difference in mean for each pair of treatments as in Eq. (6).

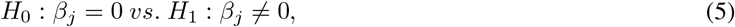

and

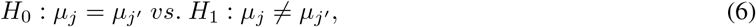

where *j* = 1, …, *m* and *j*′ = 1, …, *m* with *j* ≠ *j*′.

Finally, I consider an ordination plot of *f* (*Y*_*ij*_)’s in a reduced two- or three-dimensional coordinate space for visual representation, where the observations are later distinguished using different colors for different treatments (e.g., pre-treatment vs. post-treatment(s), or reference vs. comparison(s)).

### 2.2 Kernel Machine Comparative Analysis for Randomized Block Designs

Here, I describe the methodological details of KernRBD. First, I formulate a score test statistic using a kernel as in Eq. (7).

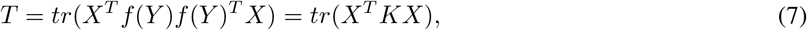

where *tr*(·) is a trace of a matrix; *K* is an *N* × *N* positive semi-definite matrix of a kernel with *K* = *f* (*Y*)*f* (*Y*)^*T*^; and *X* is an *N* × *m* design matrix of treatment labels: *X* = (*X*_11_, …, *X*_1*m*_, *X*_21_ …, *X*_2*m*_, …, *X*_*n*1_, …, *X*_*nm*_)^*T*^, where *X*_*ij*_ is an *m* × 1 vector of *m* binary (0 or 1) indicators for the subject *i* and the treatment *j*. Then, *T* is an estimate on the sum of squared overall means, subject effects and treatment effects across all subjects and treatments: 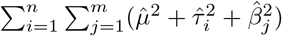.

To obtain a null test statistic value for the global testing of Eq. (3) or Eq. (4), I flip the treatment labels within each subject *i*. This is the within-block rearrangement [21], with which I switch all the rows of *X* within each subject *i*. Then, a *P*-value is calculated as in Eq. (8).

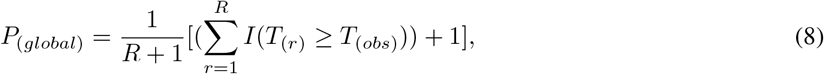

where *T*_(*r*)_ is a null test statistic value; *T*_(*obs*)_ is the observed test statistic value; and *R* is the number of possible rearrangements.

To obtain a null test statistic value for the pairwise comparison of Eq. (5) and Eq. (6), I switch only the *j*-th and *j*′-th rows of *X* within each subject *i*. Then, a *P*-value is calculated as in Eq. (9).

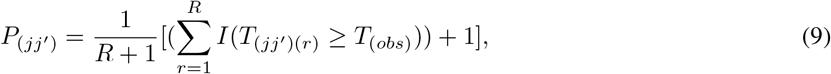

where *T*_(*jj*_*′*)(*r*) is a null test statistic value; *T*_(*obs*)_ is the observed test statistic value; *R* is the number of possible rearrangements; and *j* = 1, …, *m* and *j*′ = 1, …, *m* with *j* ≠ *j*′.

However, there are *m*!*/*(2!(*m* − 2)!) pairwise comparisons; hence, the family-wise error rate [22] cannot be controlled at a given significance level for more than two treatments (*m >* 2). As a remedy, I use the step-down procedures of [23, 24] to calculate adjusted *P*-values. For this, without loss of generality, we can suppose that there are *q* ordered pairwise comparisons (*l* = 1, …, *q*), where *l* is the order in *P*-value such that *P*_1_ ≤ · · · ≤ *P*_*q*_. Then, the adjusted *P*-values (denoted as 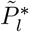) are calculated as in Eq. (10).

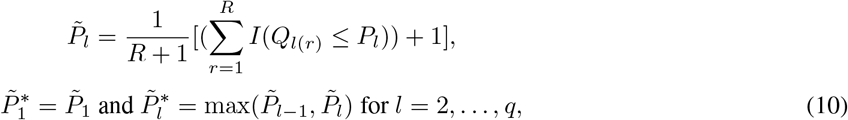

where *Q*_*q*(*r*)_ = *P*_*q*(*r*)_; *Q*_*l*(*r*)_ = min(*Q*_*l*+1(*r*)_, *P*_*l*(*r*)_) for *l* = 1, …, *q* − 1; and 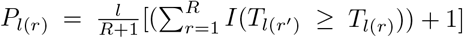 with *r*≠ *r*′.

Notably, the Mercer’s theorem [5] ensures that even when *f* (*Y*) is unknown, we can obtain its lower-dimensional representations through the singular value decomposition of a kernel matrix, *K*, as in Eq. (11).

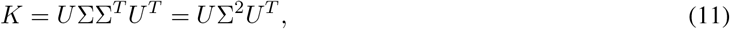

where *U* is an *N* × *N* matrix of the left eigenvectors; and Σ is an *N* × *N* diagonal matrix of the ordered eigenvalues: Σ = *diag*(*λ*_1_, …, *λ*_*N*_). Then, the columns of *U* Σ are principal components for the orthogonal and ordinal mapping of *f* (*Y*). We can pick the first two or three principal components for two- or three-dimensional visualization [17, 18, 19], wherein their proportions of total variance explained are 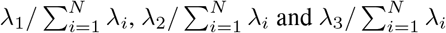.

### 2.3 Omnibus Test

Here, I describe the omnibus test of KernRBD based on the minimum *P*-value test statistic [8] as in Eq. (12) and Eq. (13) for a unified and powerful significance testing across multiple input kernels.

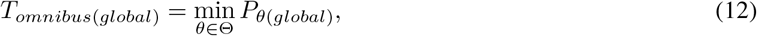

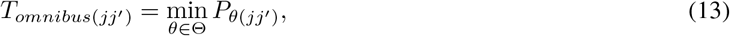

where *θ* is an index for a kernel in Θ; *P*_*θ*(*global*)_ is the *P*-value obtained through Eq. (8) based on a kernel *θ* for the global testing of Eq. (3) or Eq. (4); and *P*_*θ*(*jj*_*′*) is the *P*-value obtained through Eq. (9) based on a kernel *θ* for the pairwise comparison of Eq. (5) or Eq. (6). These minimum *P*-value test statistics reflect only the strongest evidence of significance (i.e., the minimum *P*-value) across multiple input kernels for the global testing and pairwise comparison, respectively. Therefore, powerful performance is ensured even in high-sparsity situations, where only few kernels exhibit strong significance, robustly adapting to various linear or nonlinear patterns of disparity.

Of course, the minimum *P*-value itself is a test statistic for the omnibus test, not a *P*-value finally obtained from the omnibus test. It is essential to recognize it as a random variable that depends on whether the null or alternative hypothesis holds. That is, to obtain observed test statistic values of *T*_*omnibus*(*global*)_ and *T*_*omnibus*(*jj*_*′*), *P*_*θ*(*global*)_’s and *P*_*θ*(*jj*_*′*)’s should be estimated using the original data; yet, to obtain null statistic values of *T*_*omnibus*(*global*)_ and *T*_*omnibus*(*jj*_*′*), *P*_*θ*(*global*)_’s and *P*_*θ*(*jj*_*′*)’s should be estimated using each rearranged data *r* = 1, …, *R*. More specifically, we can obtain the null test statistic values as in Eq. (14) and Eq. (15).

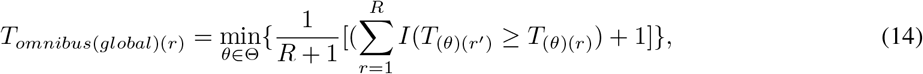

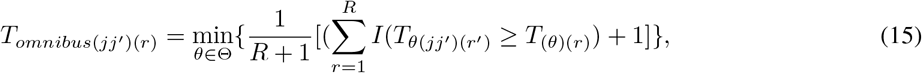

where *θ* is an index for a kernel in Θ; *j* = 1, …, *m* and *j*′ = 1, …, *m* with *j* ≠ *j*′; and *r* = 1, …, *R* and *r*′ = 1, …, *R* with *r* ≠ *r*′. Then, the *P*-values for omnibus testing are calculated as in Eq. (16) for global testing and Eq. (17) for pairwise comparisons.

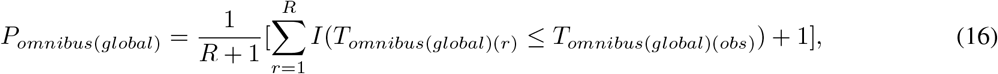

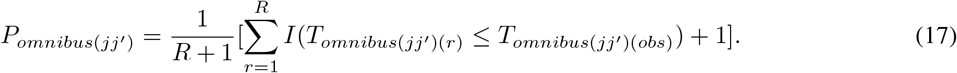

The adjusted *P*-values for more than two treatments (*m >* 2) are calculated using the same step-down procedures of [23, 24] as in Eq. (10).

## 3 Results

### 3.1 Simulation Settings

As in [4, 25], I conducted simulation experiments based on a parametric bootstrap method using the Dirichlet-multinomial distribution [26]. I generated microbial counts for the number of subjects to be 25 (*n* = 25) and 50 (*n* = 50), respectively, and the number of treatments per subject to be 4 (*m* = 4), resulting in the total numbers of observations to be 100 (*N* = 100) and 200 (*N* = 200), respectively, and the total count per observation to be 10,000. For the parameter settings, I estimated the mean proportions, 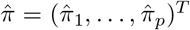, and overdispersion of the Dirichlet-multinomial distribution [26] using the Charlson et al’s upper-respiratory-tract microbiome data [27] to reflect the characteristics of real microbiome composition, including high skewness, zero-inflation and overdispersion.

To evaluate type I error for global testing and family-wise error rates for pairwise comparisons, I set the proportion parameters, *π*_*ij*_ = (*π*_*ij*1_, …, *π*_*ijp*_)^*T*^, as in Eq. (18) with the adjustment of 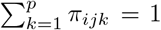 for the compositional constraint.

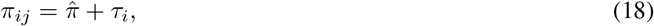

where *τ*_*i*_ is a *p* × 1 vector of the *i*-th subject’s effects: *τ*_*i*_ = (*τ*_*i*1_, …, *τ*_*ip*_)^*T*^ generated as 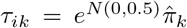 for *k* = 1, …, *p*.

To evaluate power values for global testing and family-wise confidence levels (i.e., 1 - family-wise error rates) for pairwise comparisons, I set the proportion parameters, *π*_*ij*_ = (*π*_*ij*1_, …, *π*_*ijp*_)^*T*^, as in Eq. (19) with the adjustment of 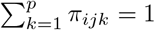 for the compositional constraint.

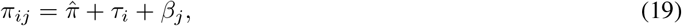

where *τ*_*i*_ is a *p* × 1 vector of the *i*-th subject’s effects: *τ*_*i*_ = (*τ*_*i*1_, …, *τ*_*ip*_)^*T*^ generated as 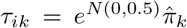 for *k* = 1, …, *p*.; and *β*_*j*_ is a *p* ×1 vector of the *j*-th treatment’s effects: *β*_*j*_ = (*β*_*j*1_, …, *β*_*jp*_)^*T*^. For the *j*-th treatment’s effects, *β*_*j*_ = (*β*_*j*1_, …, *β*_*jp*_)^*T*^, I set the microbial features in one phylogenetic cluster among ten phylogenetic clusters [28, 29] (denoted as *A*_*C*_, *C* = 1, …, 10) for the second, third and fourth treatments to have treatment effects as 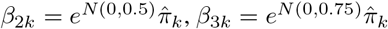 and 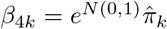, respectively, with the adjustments not to affect the other clusters, 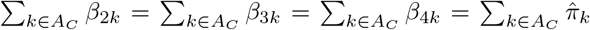, for *k* ∈ *A*_*C*_, *C* = 1, …, 10. This simulation experiment mimics various patterns of disparity with different degrees of microbial abundance, phylogenetic relatedness, cluster size, and treatment effect.

### 3.2 Simulation Results

The simulation results are organized as follows. Table 1 presents type 1 error rates and family-wise error rates for KernRBD, and type 1 error rates for PERMANOVA. Figure 1 presents power values and family-wise confidence levels for KernRBD, and power values for PERMANOVA. In Supplementary Data, S1 Table (*N* = 100) and S2 Table (*N* = 200) also present power values and family-wise confidence levels for KernRBD, and power values for PERMANOVA, using numbers.

**Table 1:** Type 1 error rates and family-wise error rates for KernRBD, and type 1 error rates for PERMANOVA (Unit: %). ^*^*K*_*J*_, *K*_*BC*_, *K*_*U*_, *K*_0.25_, *K*_0.5_, *K*_0.75_ and *K*_*W*_ represent the Jaccard, Bray-Curtis, unweighted UniFrac, generalized UniFrac (0.25), generalized UniFrac (0.5), generalized UniFrac (0.75) and weighted UniFrac kernels/distances, respec- tively. Omnibus represents the omnibus test based on the minimum *P*-value test statistic. FWER represents family-wise error rate.

| $N = 100$ | | | | | | | | | |
| --- | --- | --- | --- | --- | --- | --- | --- | --- | --- |
| | Method | $K_J$ | $K_{BC}$ | $K_U$ | $K_{0.25}$ | $K_{0.5}$ | $K_{0.75}$ | $K_W$ | Omnibus |
| Type 1 Error Rate | KernRBD | 5.01 | 4.89 | 4.97 | 5.12 | 4.92 | 5.08 | 4.92 | 4.98 |
| Type 1 Error Rate | PERMANOVA | 5.00 | 4.91 | 5.01 | 5.03 | 5.04 | 4.82 | 4.99 | - |
| FWER | KernRBD | 5.05 | 4.98 | 4.97 | 5.01 | 5.04 | 5.07 | 4.92 | 4.99 |
| FWER | PERMANOVA | - | - | - | - | - | - | - | - |
| $N = 200$ | | | | | | | | | |
| | Method | $K_J$ | $K_{BC}$ | $K_U$ | $K_{0.25}$ | $K_{0.5}$ | $K_{0.75}$ | $K_W$ | Omnibus |
| Type 1 Error Rate | KernRBD | 5.05 | 5.03 | 4.98 | 4.97 | 4.97 | 5.13 | 5.08 | 5.07 |
| Type 1 Error Rate | PERMANOVA | 5.01 | 4.85 | 5.02 | 4.97 | 5.03 | 5.14 | 5.04 | - |
| FWER | KernRBD | 5.02 | 4.84 | 4.87 | 5.17 | 4.92 | 4.96 | 5.12 | 4.92 |
| FWER | PERMANOVA | - | - | - | - | - | - | - | - |

**Figure 1:**
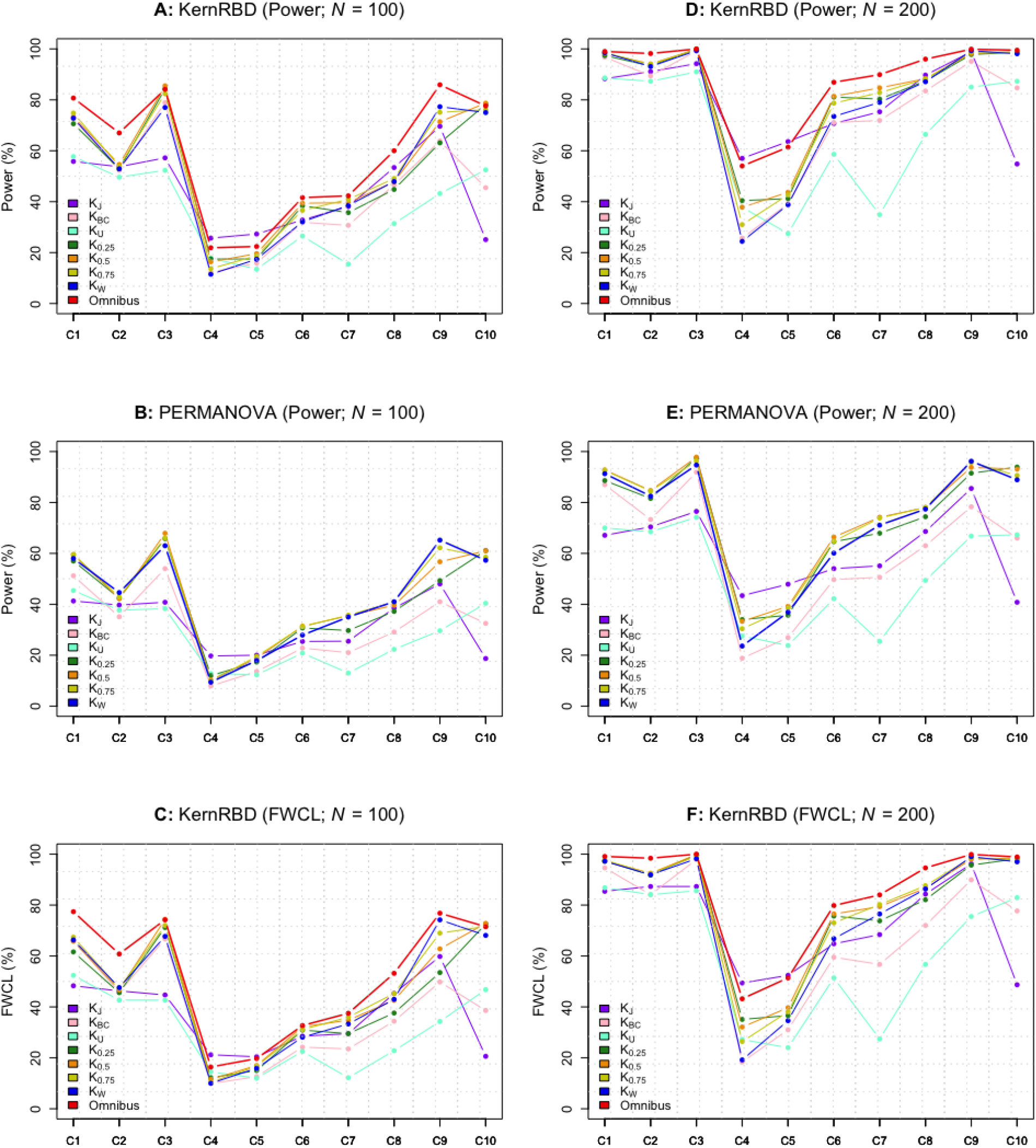
Power values and family-wise confidence levels for KernRBD, and power values for PERMANOVA (Unit: %). ^*^*K*_*J*_, *K*_*BC*_, *K*_*U*_, *K*_0.25_, *K*_0.5_, *K*_0.75_ and *K*_*W*_ represent the Jaccard, Bray-Curtis, unweighted UniFrac, generalized UniFrac (0.25), generalized UniFrac (0.5), generalized UniFrac (0.75) and weighted UniFrac kernels/distances, respec- tively. Omnibus represents the omnibus test based on the minimum *P*-value test statistic. FWCL represents family-wise confidence level. C1, …, C10 represent each phylogenetic cluster among ten phylogenetic clusters. **A** represents power values for KernRBD (*N* = 100); **B** represents power values for PERMANOVA (*N* = 100); **C** represents family-wise confidence levels for KernRBD (*N* = 100); **D** represents power values for KernRBD (*N* = 200); **E** represents power values for PERMANOVA (*N* = 200); **F** represents family-wise confidence levels for KernRBD (*N* = 200).

Note again that PERMANOVA is designed only for global testing, not providing adjusted *P*-values for pairwise comparisons, with no omnibus testing capabilities. Therefore, for PERMANOVA, only the type 1 error rates and power values for each distance are presented.

#### 3.2.1 Type 1 Error Rate and Family-Wise Error Rate

For KernRBD, we can observe well-controlled type 1 error and family-wise error rates at the 5% significance level for each kernel and the omnibus test for both *N* = 100 and *N* = 200 (Table 1). For PERMANOVA, we can observe well-controlled type 1 error rates at the 5% significance level for each distance (Table 1).

#### 3.2.2 Power and Family-Wise Confidence Level

For KernRBD, we can observe substantial variations in power values [Figure 1A and S1A Table (*N* = 100); Figure 1D and S2A Table (*N* = 200)] and family-wise confidence levels [Figure 1C and S1C Table (*N* = 100); Figure 1F and S2C Table (*N* = 200)] across different kernels and phylogenetic clusters. This shows that the performance is highly influenced by the choice of kernel and the underlying disparity pattern. However, for the omnibus test, we can observe high power values [Figure 1A and S1A Table (*N* = 100); Figure 1D and S2A Table (*N* = 200)] and high family-wise confidence levels [Figure 1C and S1C Table (*N* = 100); Figure 1F and S2C Table (*N* = 200)] across all phylogenetic clusters. This shows that the omnibus test robustly adapts to various disparity patterns.

For PERMANOVA, we can also observe substantial variations in power values [Figure 1B and S1B Table (*N* = 100); Figure 1E and S2B Table (*N* = 200)] across different distances and phylogenetic clusters; yet, PERMANOVA lacks omnibus testing capabilities. Furthermore, we can observe higher power values for KernRBD compared to PERMONOVA even for the same kernel/distance [Figure 1A > Figure 1B and S1A Table > S1B Table (*N* = 100); Figure 1D > Figure 1E and S2A Table > S2B Table (*N* = 200)].

### 3.3 Real Data Applications

Here, I demonstrate the application of KernRBD through the reanalysis of public gut microbiome data [20]. While the original data are comprehensive motivating diverse research opportunities [20], I evaluate the effect of periodically restricted feeding (PRF) on gut microbiome profiles. Specifically, the data that I use consist of 4 treatments (*m* = 4) assigned to each of 12 rhesus monkeys (*n* = 12) for 386 microbial features (*p* = 386). The first treatment serves as a baseline with no PRF intervention (denoted as Baseline), while the second (denoted as PRF 1), third (denoted as PRF 2) and fourth (denoted as PRF 3) treatments represent PRF interventions administered in a sequential order.

We can observe significant results for global testing at the 5% significance level for each kernel and the omnibus test (Table 2). However, we can observe high variability in adjusted *P*-value for pairwise comparisons (Table 2). For example, the Jaccard and uweighted UniFrac kernels yield significant results for all pairwise comparisons, though the generalized UniFrac (0.75) and weighted UniFrac kernels yield nonsignificant results for the pairwise comparisons of Baseline vs. PRF 1, Baseline vs. PRF 2, Baseline vs. PRF 3, and PRF 1 vs. PRF 2 (Table 2). Accordingly, the ordination plots show a clearer distinction across all treatments for the Jaccard (Figure 2A) and unweighted UniFrac (Figure 2C) kernels, compared to the generalized UniFrac (0.75) (Figure 2F) and weighted UniFrac (Figure 2G) kernels. Nevertheless, the omnibus test yields significant results for all pairwise comparisons (Table 2), demonstrating its strong robustness and adaptability.

**Table 2:** *P*-values for global testing and adjusted *P*-values for pairwise comparisons, organized by each kernel and the omnibus test, for KernRBD. ^*^*K*_*J*_, *K*_*BC*_, *K*_*U*_, *K*_0.25_, *K*_0.5_, *K*_0.75_ and *K*_*W*_ represent the Jaccard, Bray-Curtis, unweighted UniFrac, generalized UniFrac (0.25), generalized UniFrac (0.5), generalized UniFrac (0.75) and weighted UniFrac kernels, respectively. Omnibus represents the omnibus test based on the minimum *P*-value test statistic.

| Method | $K_J$ | $K_{BC}$ | $K_U$ | $K_{0.25}$ | $K_{0.5}$ | $K_{0.75}$ | $K_W$ | Omnibus |
| --- | --- | --- | --- | --- | --- | --- | --- | --- |
| Global Testing | <.001 | <.001 | <.001 | <.001 | <.001 | <.001 | 0.001 | <.001 |
| Baseline vs. PRF 1 | 0.006 | 0.076 | 0.028 | 0.058 | 0.172 | 0.295 | 0.365 | 0.012 |
| Baseline vs. PRF 2 | <.001 | 0.003 | <.001 | <.001 | 0.011 | 0.087 | 0.186 | <.001 |
| Baseline vs. PRF 3 | 0.003 | 0.076 | 0.010 | 0.028 | 0.056 | 0.123 | 0.186 | 0.006 |
| PRF 1 vs. PRF 2 | 0.002 | <.001 | <.001 | <.001 | 0.015 | 0.123 | 0.229 | <.001 |
| PRF 1 vs. PRF 3 | 0.006 | 0.007 | 0.004 | <.001 | <.001 | <.001 | 0.003 | <.001 |
| PRF 2 vs. PRF 3 | <.001 | <.001 | <.001 | <.001 | <.001 | 0.004 | 0.028 | <.001 |

**Figure 2:**
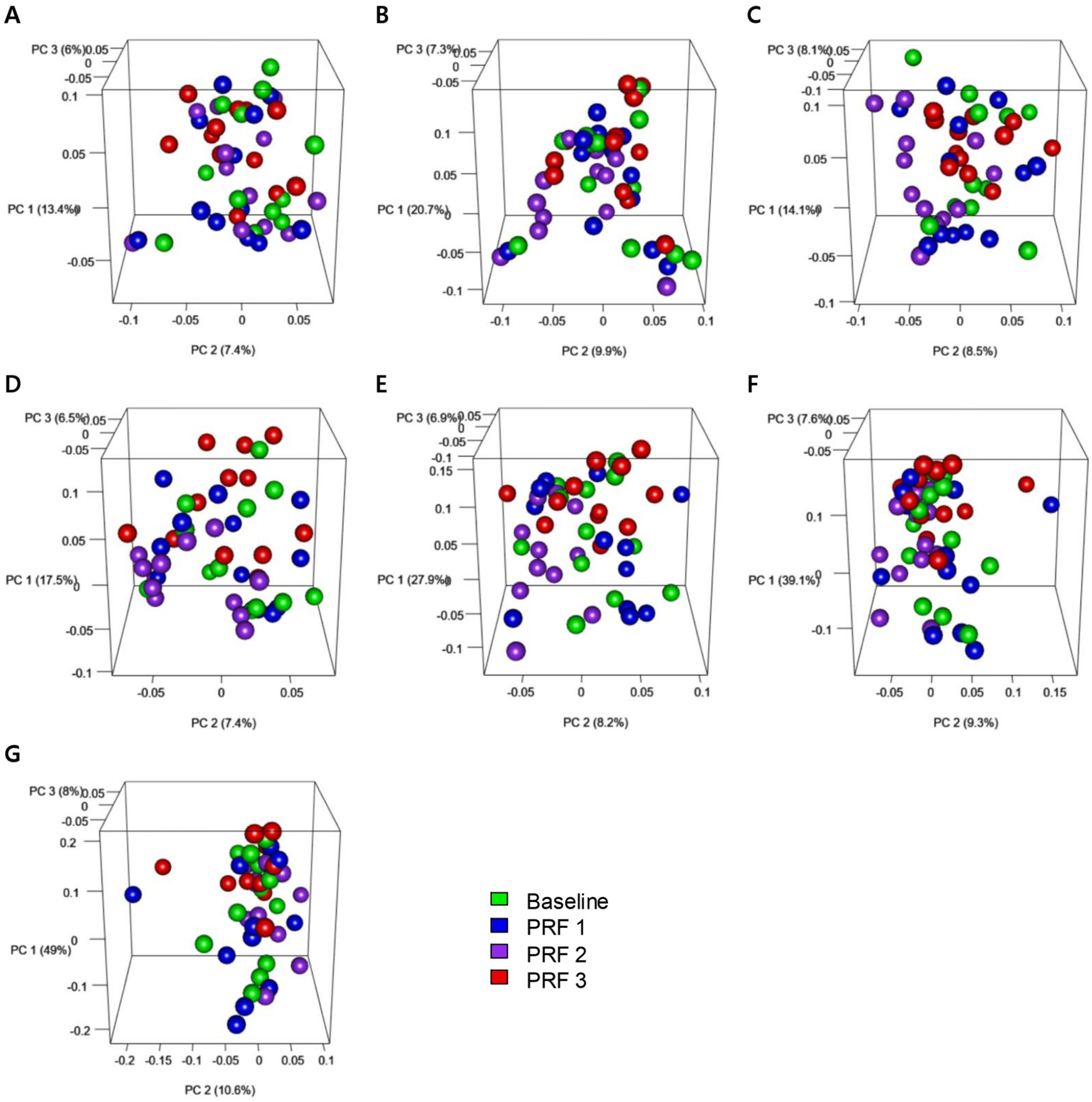
The ordination plots on the effect of PRF on gut microbiome profiles. ^*^**A, B, C, D, E, F** and **G** represent the use of Jaccard, Bray-Curtis, unweighted UniFrac, generalized UniFrac (0.25), generalized UniFrac (0.5), generalized UniFrac (0.75) and weighted UniFrac kernels, respectively.

## 4 Discussion

In this paper, I introduced KernRBD, a versatile kernel machine comparative analysis framework for randomized block designs, to assess the effects of treatments (e.g., medical interventions, environmental exposures) on high- dimensional features. KernRBD integrates the computation of *P*-value for global testing and adjusted *P*-values for pairwise comparisons, as well as the visual representation through ordination plotting. Through simulation experiments, I demonstrated that (1) KernRBD is robustly valid with well-controlled type 1 error and family-wise error rates, based on a resampling scheme that does not require the assumption of normality; (2) KernRBD outperforms PERMANOVA [9, 10, 11] in global testing, even when using the same kernel/distance; and (3) the omnibus test of KernRBD is robustly powerful across various disparity patterns. I also demonstrated the application of KernRBD through the reanalysis of public gut microbiome data to assess the effect of PRF on gut microbiome profiles [20].

I utilized ecological kernels, including Jaccard [12], Bray-Curtis [13], unweighted UniFrac [14], weighted UniFrac [15] and generalized UniFrac [16] kernels because of their unique characteristics that are well-suited for human microbiome *β*-diversity analysis. However, I do not limit the application of KernRBD to these ecological kernels or to human microbiome *β*-diversity analysis. KernRBD is a practical tool that requires only a kernel as input, even without the need to understand its underlying feature transformations. In practice, there are many other kernels, and the suitable set of candidate input kernels can vary depending on the field of study. Developing new kernels can also be needed to deepen our understanding of various omics profiles, and hence to fully optimize the use of KernRBD. However, I was unable to address all of these possibilities in this study.

## Data Availability

I utilized public microbiome data to assess the effect of PRF on gut microbiome profiles [20]. The raw sequence data are deposited in NCBI Gene Expression Omnibus (http://www.ncbi.nlm.nih.gov/geo; accession number GSE235769). The processed data are available as example data to implement KernRBD in the R package, KernRBD (https://github.com/hk1785/kernrbd).

