## Supplementary Data for "A general kernel machine comparative analysis framework for randomized block designs"

S1 Table: Power values and family-wise confidence levels for KernRBD, and power values for PERMANOVA for $N = 100$ (Unit: %). *$K_J$, $K_{BC}$, $K_U$, $K_{0.25}$, $K_{0.5}$, $K_{0.75}$ and $K_W$ represent the Jaccard, Bray-Curtis, unweighted UniFrac, generalized UniFrac (0.25), generalized UniFrac (0.5), generalized UniFrac (0.75) and weighted UniFrac kernels/distances, respectively. Omnibus represents the omnibus test based on the minimum *P*-value test statistic. FWCL represents family-wise confidence level. C1, . . ., C10 represent each phylogenetic cluster among ten phylogenetic clusters. **A** represents power values for KernRBD ($N = 100$); **B** represents power values for PERMANOVA ($N = 100$); **C** represents family-wise confidence levels for KernRBD ($N = 100$).

| A: KernRBD (Power; $N = 100$) | | | | | | | | | | |
|---|---|---|---|---|---|---|---|---|---|---|
| | C1 | C2 | C3 | C4 | C5 | C6 | C7 | C8 | C9 | C10 |
| $K_J$ | 55.8 | 53.7 | 57.2 | 25.7 | 27.3 | 32.8 | 38.1 | 53.4 | 69.6 | 25.1 |
| $K_{BC}$ | 72.6 | 52.3 | 78.9 | 12.2 | 15.9 | 31.9 | 30.7 | 46.6 | 63.8 | 45.5 |
| $K_U$ | 57.7 | 49.6 | 52.3 | 17.9 | 13.5 | 26.5 | 15.5 | 31.4 | 43.2 | 52.5 |
| $K_{0.25}$ | 70.6 | 52.9 | 84.3 | 17.5 | 17.5 | 38.4 | 35.7 | 44.8 | 63.1 | 78.2 |
| $K_{0.5}$ | 73.3 | 54.6 | 85.5 | 16.4 | 19.6 | 39.4 | 39.6 | 47.8 | 71.4 | 78.7 |
| $K_{0.75}$ | 74.8 | 53.8 | 82.4 | 13.7 | 18.8 | 36.6 | 40.9 | 49.1 | 75.1 | 76.1 |
| $K_W$ | 72.8 | 52.9 | 77.0 | 11.5 | 17.4 | 32.1 | 38.5 | 47.9 | 77.3 | 75.0 |
| Omnibus | 80.7 | 67.0 | 84.1 | 21.9 | 22.4 | 41.6 | 42.3 | 60.0 | 85.9 | 77.6 |

| B: PERMANOVA (Power; $N = 100$) | | | | | | | | | | |
|---|---|---|---|---|---|---|---|---|---|---|
| | C1 | C2 | C3 | C4 | C5 | C6 | C7 | C8 | C9 | C10 |
| $K_J$ | 41.3 | 39.7 | 40.8 | 19.7 | 20.0 | 25.4 | 25.5 | 38.5 | 48.0 | 18.7 |
| $K_{BC}$ | 51.2 | 35.1 | 54.0 | 7.9 | 13.6 | 22.8 | 21.0 | 29.1 | 41.0 | 32.5 |
| $K_U$ | 45.4 | 37.7 | 38.3 | 12.8 | 12.3 | 20.8 | 13.0 | 22.3 | 29.6 | 40.4 |
| $K_{0.25}$ | 57.0 | 42.4 | 65.8 | 12.0 | 17.3 | 30.7 | 29.7 | 37.2 | 49.3 | 61.1 |
| $K_{0.5}$ | 59.5 | 42.2 | 67.9 | 10.3 | 18.8 | 31.4 | 35.5 | 39.6 | 56.7 | 61.0 |
| $K_{0.75}$ | 59.6 | 42.8 | 66.1 | 9.8 | 19.5 | 31.2 | 35.7 | 40.9 | 62.2 | 58.5 |
| $K_W$ | 58.0 | 44.6 | 63.0 | 9.4 | 17.9 | 27.9 | 35.0 | 41.0 | 65.2 | 57.3 |
| Omnibus | - | - | - | - | - | - | - | - | - | - |

| C: KernRBD (FWCL; $N = 100$) | | | | | | | | | | |
|---|---|---|---|---|---|---|---|---|---|---|
| | C1 | C2 | C3 | C4 | C5 | C6 | C7 | C8 | C9 | C10 |
| $K_J$ | 48.3 | 46.3 | 44.7 | 21.2 | 20.4 | 28.5 | 29.5 | 45.3 | 59.8 | 20.6 |
| $K_{BC}$ | 66.0 | 45.8 | 66.7 | 9.9 | 12.6 | 24.2 | 23.5 | 34.4 | 49.8 | 38.6 |
| $K_U$ | 52.4 | 42.7 | 42.7 | 14.5 | 12.0 | 22.5 | 12.2 | 22.8 | 34.3 | 46.8 |
| $K_{0.25}$ | 61.6 | 45.6 | 71.2 | 12.2 | 15.0 | 30.9 | 29.5 | 37.6 | 53.5 | 72.6 |
| $K_{0.5}$ | 65.8 | 46.7 | 74.4 | 11.5 | 17.0 | 32.2 | 34.8 | 42.6 | 62.8 | 72.8 |
| $K_{0.75}$ | 67.4 | 47.3 | 72.0 | 10.4 | 16.7 | 31.1 | 36.0 | 45.3 | 69.0 | 71.6 |
| $K_W$ | 66.3 | 47.6 | 67.7 | 10.0 | 15.9 | 28.1 | 33.3 | 43.0 | 74.2 | 68.1 |
| Omnibus | 77.4 | 60.8 | 74.2 | 16.4 | 19.7 | 32.7 | 37.5 | 53.2 | 76.8 | 71.5 |

S2 Table: Power values and family-wise confidence levels for KernRBD, and power values for PERMANOVA for $N = 200$ (Unit: %). *$K_J$, $K_{BC}$, $K_U$, $K_{0.25}$, $K_{0.5}$, $K_{0.75}$ and $K_W$ represent the Jaccard, Bray-Curtis, unweighted UniFrac, generalized UniFrac (0.25), generalized UniFrac (0.5), generalized UniFrac (0.75) and weighted UniFrac kernels/distances, respectively. Omnibus represents the omnibus test based on the minimum *P*-value test statistic. FWCL represents family-wise confidence level. C1, ..., C10 represent each phylogenetic cluster among ten phylogenetic clusters. **A** represents power values for KernRBD ($N = 200$); **B** represents power values for PERMANOVA ($N = 200$); **C** represents family-wise confidence levels for KernRBD ($N = 200$).

| A: KernRBD (Power; $N = 200$) | | | | | | | | | | |
|---|---|---|---|---|---|---|---|---|---|---|
| | C1 | C2 | C3 | C4 | C5 | C6 | C7 | C8 | C9 | C10 |
| $K_J$ | 88.3 | 91.1 | 94.2 | 57.0 | 63.6 | 70.7 | 75.3 | 89.7 | 98.9 | 54.8 |
| $K_{BC}$ | 96.8 | 89.4 | 99.2 | 25.7 | 39.2 | 71.1 | 71.9 | 83.5 | 95.1 | 84.7 |
| $K_U$ | 88.6 | 87.3 | 91.0 | 37.8 | 27.5 | 58.6 | 34.9 | 66.4 | 85.0 | 87.3 |
| $K_{0.25}$ | 97.4 | 93.3 | 99.9 | 40.4 | 41.2 | 81.1 | 80.3 | 87.4 | 97.8 | 98.7 |
| $K_{0.5}$ | 98.3 | 94.1 | 99.9 | 37.8 | 43.7 | 81.4 | 84.7 | 88.3 | 98.2 | 98.9 |
| $K_{0.75}$ | 98.5 | 93.9 | 99.8 | 31.0 | 42.7 | 78.7 | 82.8 | 88.2 | 98.7 | 98.6 |
| $K_W$ | 98.5 | 93.1 | 99.3 | 24.4 | 38.8 | 73.5 | 79.0 | 87.1 | 99.3 | 98.1 |
| Omnibus | 99.0 | 98.2 | 99.9 | 54.0 | 61.4 | 86.9 | 89.9 | 96.0 | 99.9 | 99.5 |
| **B: PERMANOVA (Power; $N = 200$)** | | | | | | | | | | |
| | C1 | C2 | C3 | C4 | C5 | C6 | C7 | C8 | C9 | C10 |
| $K_J$ | 67.1 | 70.4 | 76.5 | 43.4 | 47.9 | 54.0 | 55.1 | 68.6 | 85.5 | 40.8 |
| $K_{BC}$ | 87.0 | 73.3 | 91.9 | 18.8 | 26.9 | 49.7 | 50.6 | 63.0 | 78.3 | 66.0 |
| $K_U$ | 70.0 | 68.5 | 74.2 | 27.5 | 23.8 | 42.2 | 25.4 | 49.4 | 66.8 | 67.3 |
| $K_{0.25}$ | 88.6 | 81.6 | 97.5 | 34.0 | 35.7 | 64.6 | 67.9 | 74.4 | 91.5 | 93.9 |
| $K_{0.5}$ | 92.8 | 84.7 | 97.8 | 33.3 | 39.2 | 66.4 | 74.2 | 78.0 | 93.8 | 93.1 |
| $K_{0.75}$ | 92.6 | 84.5 | 96.3 | 30.4 | 38.5 | 64.8 | 73.9 | 78.1 | 95.7 | 90.6 |
| $K_W$ | 91.3 | 82.5 | 94.7 | 23.6 | 36.9 | 60.1 | 71.1 | 77.4 | 96.2 | 88.9 |
| Omnibus | - | - | - | - | - | - | - | - | - | - |
| **C: KernRBD (FWCL; $N = 200$)** | | | | | | | | | | |
| | C1 | C2 | C3 | C4 | C5 | C6 | C7 | C8 | C9 | C10 |
| $K_J$ | 85.4 | 87.3 | 87.3 | 49.4 | 52.4 | 64.8 | 68.4 | 84.3 | 96.2 | 48.7 |
| $K_{BC}$ | 94.6 | 84.1 | 97.9 | 18.4 | 31.0 | 59.5 | 56.7 | 72.0 | 89.9 | 77.7 |
| $K_U$ | 86.8 | 84.1 | 85.6 | 27.2 | 24.0 | 51.4 | 27.4 | 56.7 | 75.5 | 82.9 |
| $K_{0.25}$ | 97.4 | 91.8 | 99.8 | 35.1 | 36.6 | 75.7 | 73.8 | 82.1 | 95.7 | 98.2 |
| $K_{0.5}$ | 97.3 | 92.5 | 99.8 | 32.1 | 39.7 | 76.5 | 79.4 | 86.6 | 98.0 | 98.5 |
| $K_{0.75}$ | 97.7 | 92.2 | 99.2 | 26.4 | 38.1 | 73.0 | 80.2 | 87.6 | 98.6 | 97.6 |
| $K_W$ | 97.2 | 91.9 | 98.2 | 19.2 | 34.6 | 66.8 | 76.5 | 86.3 | 99.0 | 97.0 |
| Omnibus | 99.1 | 98.4 | 99.9 | 43.2 | 51.4 | 79.8 | 84.0 | 94.6 | 99.9 | 98.9 |
